# Signatures of Environmental Adaptation During Range Expansion of Wild Common Bean (*Phaseolus vulgaris*)

**DOI:** 10.1101/571042

**Authors:** Andrea Ariani, Paul Gepts

## Abstract

Landscape genomics integrates population genetics with landscape ecology, allowing the identification of putative molecular determinants involved in environmental adaptation across the natural geographic and ecological range of populations. Wild *Phaseolus vulgaris*, the progenitor of common bean (*P. vulgaris*), has a remarkably extended distribution over 10,000 km from northern Mexico to northwestern Argentina. Earlier research has shown that this distribution represents a range expansion from Mesoamerica to the southern Andes through several discrete migration events and that the species colonized areas with different temperature and rainfall compared to its core area of origin. Thus, this species provides an opportunity to examine to what extent adaptation of a species can be broadened or, conversely, ecological or geographical distribution can be limited by inherent adaptedness. In the current study, we applied a landscape genomics approach to a collection of 246 wild common bean accessions representative of its broad geographical and climatic distribution and genotyped for ∼20K SNPs. We applied two different but complementary approaches for identifying loci putatively involved in environmental adaptation: i) an outlier-detection method that identifies loci showing strong differentiation between sub-populations; ii) an association method based on the identification of loci associated with bio-climatic variables. This integrated approach allowed the identification of several genes showing signature of selection across the different natural sub-populations of this species, as well as genes associated with specific bio-climatic variables related to temperature and precipitation. The current study demonstrates the feasibility of landscape genomics approach for a preliminary identification of specific populations and novel candidate genes involved in environmental adaptation in *P. vulgaris*. As a resource for broadening the genetic diversity of the domesticated gene pool of this species, the genes identified constitute potential molecular markers and introgression targets for the breeding improvement of domesticated common bean.

**Author Summary:** The ancestral form of common bean has an unusually large distribution in the Americas, extending over 10,000 km from ∼35° N. Lat. to ∼35° S. Lat. This wide distribution results from discrete long-range dissemination events to the Andes region from the original environments in Mesoamerica. It also suggests adaptation to new environments that are distinct from those encountered in Mesoamerica. In this research, we identified genes that may be involved in adaptation to climate variables in these new environments using two methods. A first method – outlier detection – was used to identify genome regions that differentiated the wild bean groups in the Andes resulting from discrete dissemination events among themselves and the different groups in Mesoamerica. The second method – genome-wide association – was used to identify candidate genome regions correlated with these same variables across the entire distribution from Mesoamerica to the southern Andes. The two methods identified two sets of candidate genes, several of which were related to the water status of plants, and illustrate how the genetic architecture of adaptation following long-range dissemination. This study provides sets of candidate genes as well as candidate wild bean populations that need to be corroborated for their use in increasing the water use efficiency of domesticated beans.

## Introduction

Climate change represents one of the primary threats for food security worldwide, but especially in developing countries that rely heavily on agricultural production from smallholder farmers (Rippke et al., 2016; Campbell et al., 2016). Indeed, several studies have highlighted a predominant role of climate change in reducing agricultural productivity and increasing inter-annual variability in crop yields, thus directly affecting food availability and stability (Wheeler and von Braun, 2013; Challinor et al., 2014). The increase in average temperatures, along with the higher frequency and intensity of extreme weather conditions, will require the development of new plant varieties adapted to this changing environment in order to meet future food security needs (Lobell et al., 2008; Field et al., 2012). The development of new varieties requires the introduction of genetic diversity into breeding programs to find the correct combinations of favorable alleles in a specific crop (Ford-Lloyd et al., 2011). The genetic variability available in domesticated plants is generally low due to the bottleneck effect induced by domestication and subsequent selection during variety improvement (Ford-Lloyd et al., 2011; Zamir 2001; Gepts 2014), thus new sources of genetic diversity need to be introduced into breeding programs.

Crop Wild Relatives (CWRs) represent a large, and mostly unexploited, source of genetic diversity readily available for plant improvement under climate change (Ford-Lloyd et al., 2011; Zamire 2001; Gepts 2014; Spillane and Gepts 2001; Brozynska et al., 2016). However, the use of CWRs in breeding programs for improving stress resistance in domesticated species could be hindered by the lack of knowledge of the genetic determinants of resistance, difficulties in phenotyping a large number of individuals under agricultural conditions, and the existence of linkages between target resistance genes and unfavorable loci subject to linkage drag (Brozynska et al., 2016; Cortés et al., 2013; Zhang et al., 2017). One possible solution for overcoming the first two difficulties is the integration of environmental and genotypic datasets to understand the genetic basis of natural selection in wild populations, an approach known as ‘landscape genomics’ (Schoville et al., 2012; Bragg et al., 2015). In addition, this approach offers both theoretical and practical applications since it strengthens the understanding of plant natural adaptation but allows also the identification of germplasm accessions and molecular markers that could be readily applicable – pending validation - for breeding improvement of domesticated plants (Anderson et al., 2016).

Several methods have been developed for identifying signatures of natural selection (e.g., selective sweeps) in natural populations. These methods can be divided mostly in outlier-detection methods, which identify hard-selection sweeps, and association methods, which identify soft-selection sweeps (Schoville et al., 2012; Wagner and Fortin, 2013). Outlier-detection methods are based on population differentiation analysis and aim at identifying loci with drastic differences in allele frequencies between populations, as measured by *F_st_* (Wright, 1949; Lewontin and Krakauer, 1973). Although based on the assumption that alleles fixed within sub-populations could confer an evolutionary advantage in the ecological niche occupied (Haldane, 1930; Kimura, 1962), these methods do not take directly into account climatic data and could be biased by complex population structure and/or demography (Narum and Hess, 2011). On the other hand, association methods directly correlate genotypic with environmental data and rely on the assumption that variations of allele frequencies across environmental gradients are possible signature of local adaptation (Manel et al., 2010). The theory beneath environmental association methods are practically the same as that used in Genome Wide Association Studies (GWAS) (Hirschhorn and Daly, 2005). Both approaches employ mixed model association approaches for correcting the confounding effects that could be introduced by population structure and relatedness in the sample (Lipka et al., 2015).

Common bean (*Phaseolus vulgaris* L.) is an essential staple crop providing most of proteins and micronutrients in the diet of the majority of the population in several developing countries (Gepts et al., 2008). The regular consumption of this crop provides several health benefits, like reducing the risks of heart disease, obesity, and diabetes (Messina, 2014). Its cultivation improves agricultural sustainability thanks to its nitrogen-fixing ability (Rubiales and Mikić, 2015). Common bean shows a surprisingly high genetic diversity, with the presence of at least three geographically isolated and divergent wild gene pools located in 1) Mesoamerica and the northern Andes (MW); 2) the Central Andes (Ecuador and northern Peru; PhI); and 3) the Southern Andes (southern Peru, Bolivia, and northwestern Argentina; AW) (Chacón et al., 2007; Koenig and Gepts, 1989; Debouck et al., 1993; Mamidi et al., 2013). Common bean was domesticated independently in Mexico and the Southern Andes, producing locally-adapted varieties and landraces with specific characteristics (Bitocchi et al., 2013; Blair et al., 2012; Gepts et al., 1986, Mamidi et al., 2011, Rossi et al., 2009, Singh et al., 1991). The intermediate gene pool in the Central Andes was not domesticated (Debouck et al., 1993; Kami et al., 1995). This wild group has been recently identified as a cryptic sister species of *P. vulgaris*, named *Phaseolus debouckii* A. Delgado, which was disseminated from the center of origin of this species in Mesoamerica and remained geographically isolated from the other wild gene pools of this species (Rendón-Anaya et al., 2017a,b).

Wild common bean is an annual vine plant, which is distributed from the state of Chihuahua in northern Mexico (approx. 35° N. Lat.) to the Córdoba province in Argentina (approx. 35° S. Lat.), encompassing almost 70 latitudinal degree or about 10,000 km (Gepts, 1998; Porch et al., 2013). This species grows in both tropical and sub-tropical environments across the Americas at elevations between 500 and 2,000 m a.s.l. with annual rainfall from 500 to 1,800 ml (Cortés et al., 2013, Gepts, 1998, Porch et al., 2013). This broad geographic and ecological distribution suggests the existence of genotypes adapted to a wide variety of environmental conditions, which could be useful donors of abiotic stress resistance for improving domesticated common bean production under climate change (Porch et al., 2013, Acosta-Gallegos et al., 2007).

Future projection of climate changes under different models predict a reduction of suitability for common bean production in areas where this plant is an essential staple crop and also a source of household income, hence endangering food security and increasing rural poverty in already susceptible areas of the world (Ramirez-Cabral et al., 2016). For this reason, it is essential to understand the molecular mechanisms involved in wild common bean adaptation to different environments and to identify molecular markers that could be useful in breeding improvement of this crop. The application of landscape genomics approaches in wild common bean could help address these issues, as demonstrated previously in several other plant species like soybean, barley, *Medicago truncatula*, maize, and *Brachypodium* (Anderson et al., 2016, Westengen et al., 2012, Yoder et al., 2014, Dell'Acqua et al., 2014, Abebe et al., 2015).

In the current study, we applied a landscape genomics approach to understand environmental adaptation to a dataset comprised of 246 wild common beans genotyped for ∼20K previously developed SNPs (Ariani et al., 2018). A similar analysis was performed previously in this species using 148 SNPs located in genes putatively involved in adaptation to biotic or abiotic stresses (Rodriguez et al., 2016). However, the higher number of markers developed in this study and the broader and more even distribution across the genome of these markers, results in a more comprehensive and precise analysis of environmental adaptation in this species. In addition, the genes identified as associated with environmental variables can be validated and applied in the future for domesticated common bean breeding improvement.

## Results

### Bio-climatic data analysis

The bio-climatic variables downloaded from the WorldClim database concern mostly temperature and rainfall during the year. These bio-climatic variables were developed for generating biologically informative variables useful for species distribution modeling and landscape genomics approaches. In our analyses, the 19 bio-climatic variables analyzed showed a great degree of correlation, in particular for similar variables like bio_14 (precipitation of the driest month) and bio_17 (precipitation of the driest quarter), or bio_13 (precipitation of the wettest month) with bio_16 (precipitation of the wettest quarter) (**S1 Fig**).

The loading plot on the first two PCs showed some correlations between bio-climatic variables and principal components, as well as strong correlations between some of the bio-climatic variables analyzed (Fig 1A). In particular, bio_12 (annual precipitation) and bio_4 (temperature seasonality) showed a strong correlation with PC1. On the other hand, bio_5 (max temperature of the warmest month), bio_8 (mean temperature of the wettest quarter), and bio_10 (mean temperature of the warmest quarter) showed a strong correlation with PC2. Interestingly, most of the variables related to precipitation (bio_12, bio_14, bio_16, bio_17, bio_18, and bio_19) were positively correlated with PC1, the variables related to seasonal variation (bio_2, bio_4, bio_7, and bio_15) were negatively correlated with PC1, while the variables related to temperature (bio_1, bio_5, bio_8, bio_9, bio_10, and bio_11) were negatively correlated with PC2.

**Fig 1.**
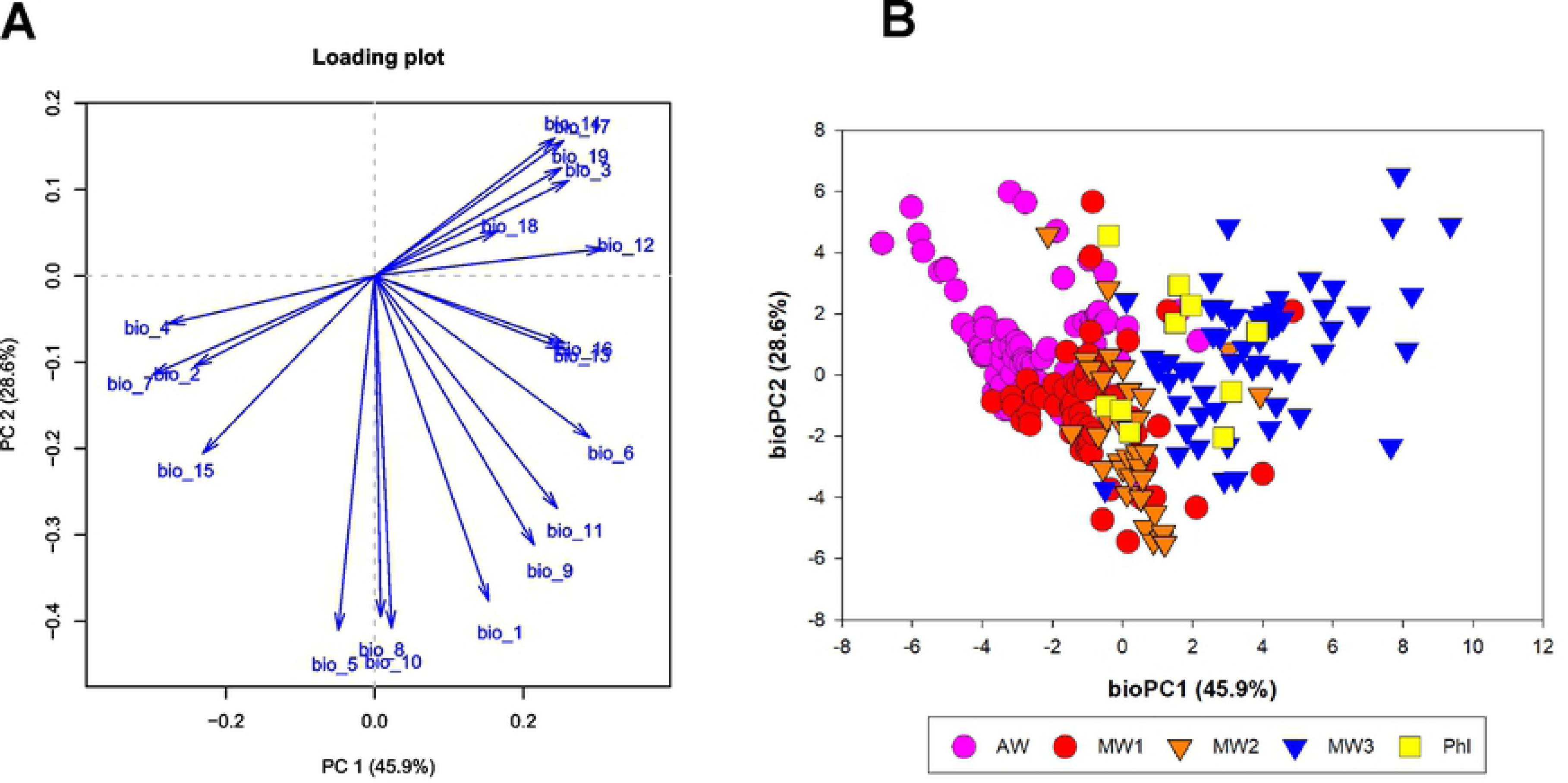
Bio-climatic data analysis. **(A)** Loading plot of the PCA analysis. **(B)** Principal Component Analysis (PCA) of the bio-climatic data. Groups are colored according to the K-mean clustering analysis conducted in this study, which gave results very similar to the STRUCTURE analysis conducted by Ariani et al. (2018): MW1, MW2, and MW3: Mesoamerican wild gene pools; AW: Andean wild gene pool; PhI: Intermediate wild gene pool.

In addition, this PCA on the bio-climatic variables for the genotypes analyzed showed that the first two principal components (PC1 and PC2) explained 75% of the variance (Fig 1B), while PC1 to PC4 explained cumulatively > 90% of the variance (**S2 Fig**). A plot of PC1 vs. PC2 showed some differences in the distribution of the different gene pools of wild common bean in the PC dimensional space. In particular, the majority of genotypes from the Mesoamerican (MW1 to MW3) and Intermediate (PhI) gene pools were distributed towards the positive part of PC1, while the Andean group were located in the negative part of this axis (Fig. 1A). Given the origin of the genus *Phaseolus* in the Mesoamerican area (with local descendants represented by MW1 and MW2), three range expansions characterize this species: 1) PhI, which established wild populations on the western slope of the Andes in Ecuador and northern Peru; 2) AW, encompassing wild populations in the southern Andes; and 3) MW3, a more recent and perhaps ongoing dissemination to Central America and the eastern slope of the northern Andes (Ariani et al., 2018). Inspection of Fig 1A and **S3 Table** shows that the distribution of the PhI group, which resulted from the earliest range expansion event, correlates - on bioPC3 - with Isothermality (bio_3), Temperature Seasonality (bio_4), bio_13 (Precipitation of the Wettest Month), and bio_18 (Precipitation of the Warmest Quarter), consistent with a dispersal to an equatorial region. In contrast, the predominant distribution of the southern Andean accessions (AW) in the upper left quadrant of Fig 1 is consistent with earlier observations that the populations of this gene pool are distributed in cooler and drier locations, as shown by correlations with bio_6 (Minimum Temperature of the Coldest Month), bio_9 (Mean Temperature of the Driest Quarter, bio_11 (Mean Temperature of the Coldest Quarter) and bio_1 (Annual Mean Temperature). This dissemination occurred with a concomitant lower potential evapotranspiration (Ariani et al. 2018). Dispersal of the MW3 group (Fig 1) increased Isothermality (bio_3) and decreased Seasonality (bio_4) and Precipitation Seasonality (bio_15); it also increased Precipitation during the Driest Month (bio_14) and Driest Quarter (bio_17).

### Genome scan of selection

An analysis of the scree plot of the PCA analysis conducted on SNP data (molecular PCA) showed that a quarter of the variance could be explained by the first principal component, even though molPC2 to molPC5 also explained a considerable amount of variance in the data (Fig 2A). On the other hand, after molPC5, no large increase in the cumulative explained variance could be detected. This pattern of the scree plot is representative of a possible range expansion of this species across the Americas, as hypothesized by a prior evolutionary analysis of this same collection (Ariani et al., 2018). Visual inspection of p-value distribution for genome scans for K=2 and K=3 showed a large proportion of low and high p-values, while for K=4 and K=5 the distribution of p-values was more uniform, especially for K=5 (**S3 Fig**). For this reason, we selected K=5 for further genome scan analysis.

**Fig 2.**
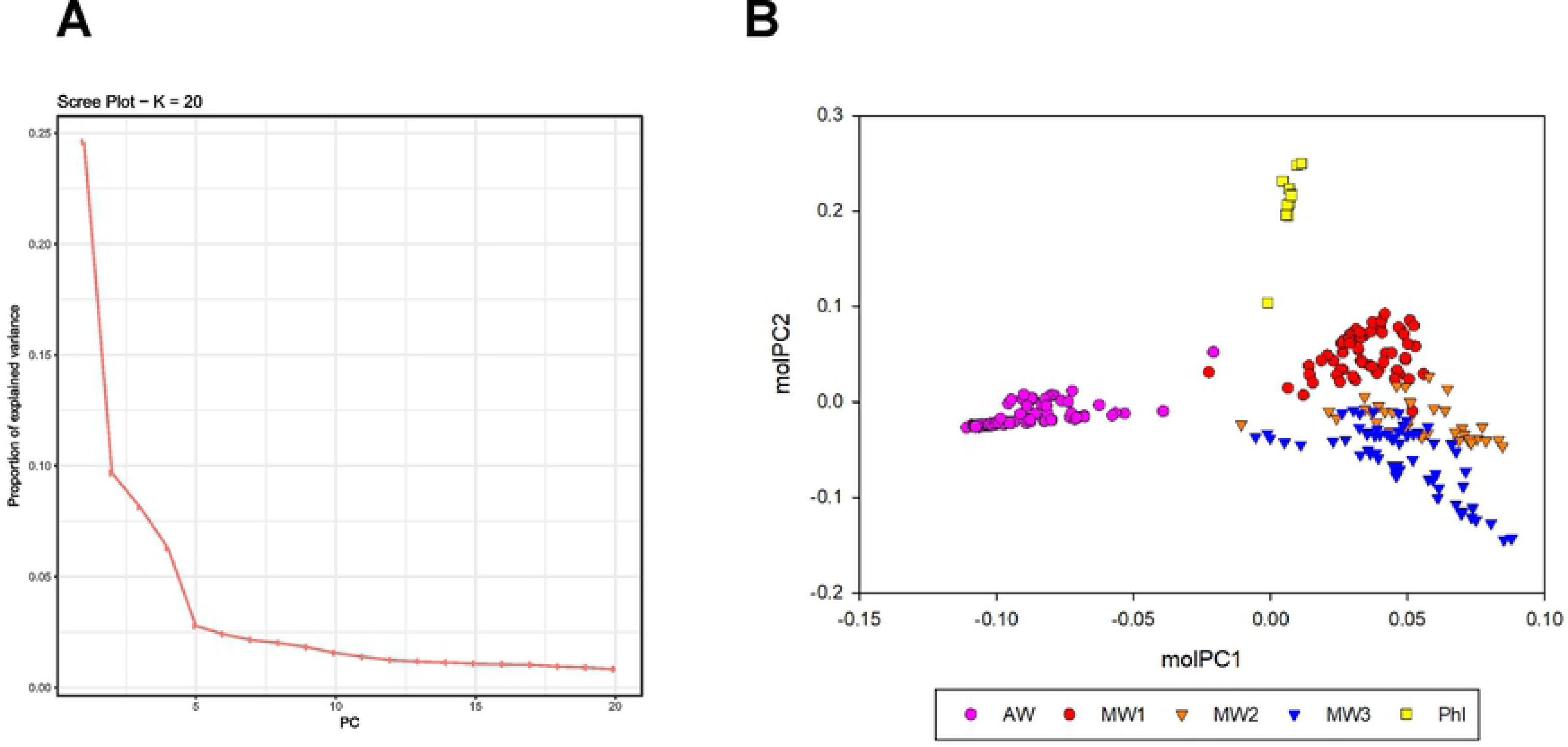
Principal Component Analysis on SNP data. **(A)** Screeplot of the PCA explained variance. **(B)** PCA plot based on molecular data of the different genotyped analyzed in the current study. Groups are colored according to the K-mean clustering analysis conducted in this study, which gave results very similar to the STRUCTURE analysis conducted by Ariani et al. (2018): MW1, MW2, and MW3: Mesoamerican wild gene pools; AW: Andean wild gene pool; PhI: Intermediate wild gene pool.

A plot of genetic PCA analysis performed with the pcadapt algorithm was able to discriminate between the different wild gene pools of this species (Fig 2B). In particular, the MW1-MW3 groups were mostly localized on the positive part of the molecular PC1 (molPC1) axis, while the AW gene pool was localized towards the negative end of molPC1. Interestingly, molPC1 mostly differentiated MW vs. AW, while molPC2 and molPC3 clearly separated the MW+AW groups from the PhI (Fig 3).

**Fig 3.**
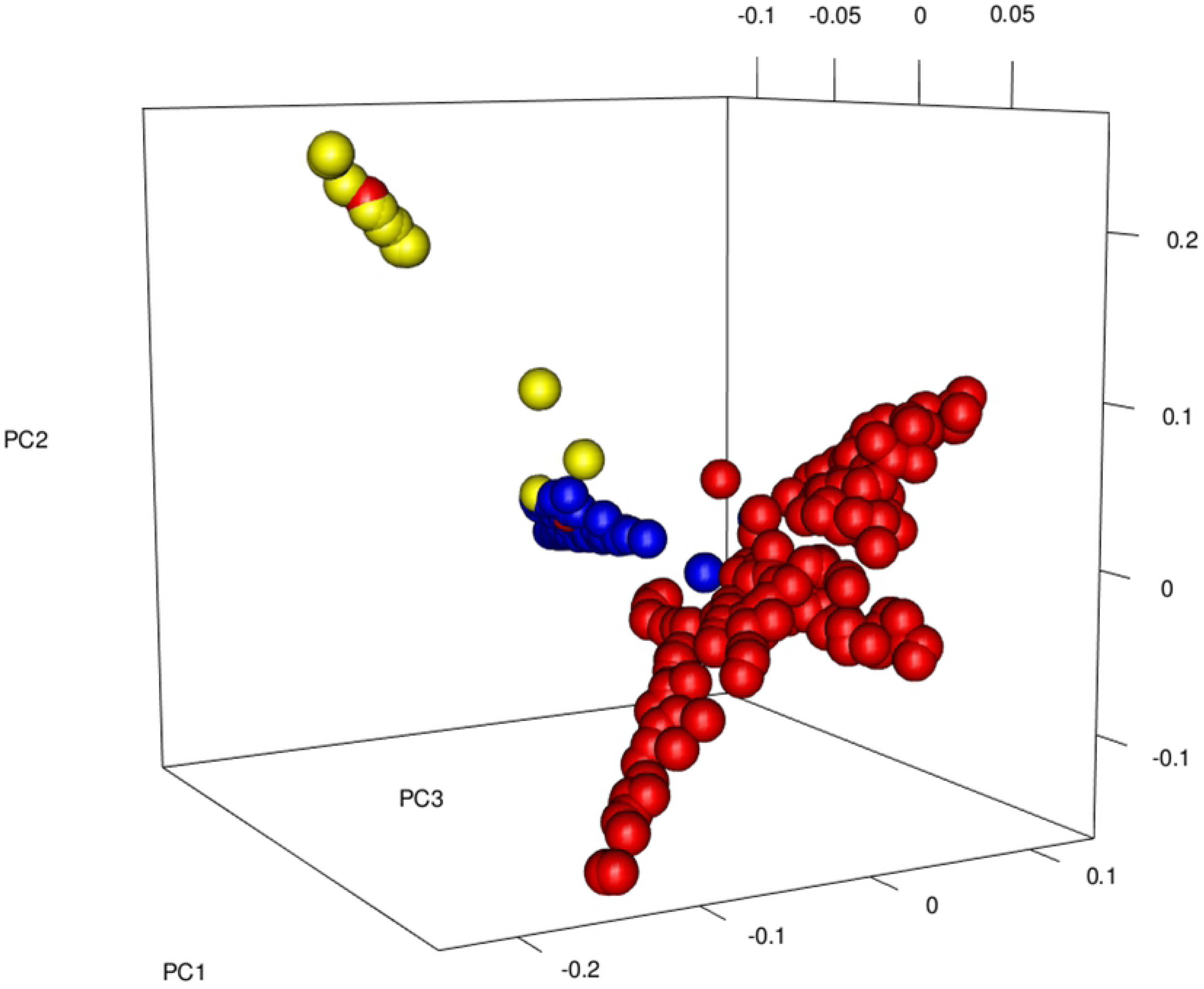
Three-dimensional plot of the PCA analysis on molecular data. Points are colored as in Fig 2B.

The genome scan analysis with K=5 identified 84 significant variants (Bonferroni-corrected p-value ≤ 0.001) distributed throughout the 11 chromosomes of common bean (Fig 4), tagging 70 annotated genes (**S1 Table**). The highest number of tagged genes were identified on chromosomes Pv02 and Pv04 with 15 and 11 genes, respectively. The genes identified as selected by genome scan analysis were mostly related to plant development (17 genes), hormone response (10 genes), ion homeostasis (5 genes), and response to stress (9 genes).

**Fig 4.**
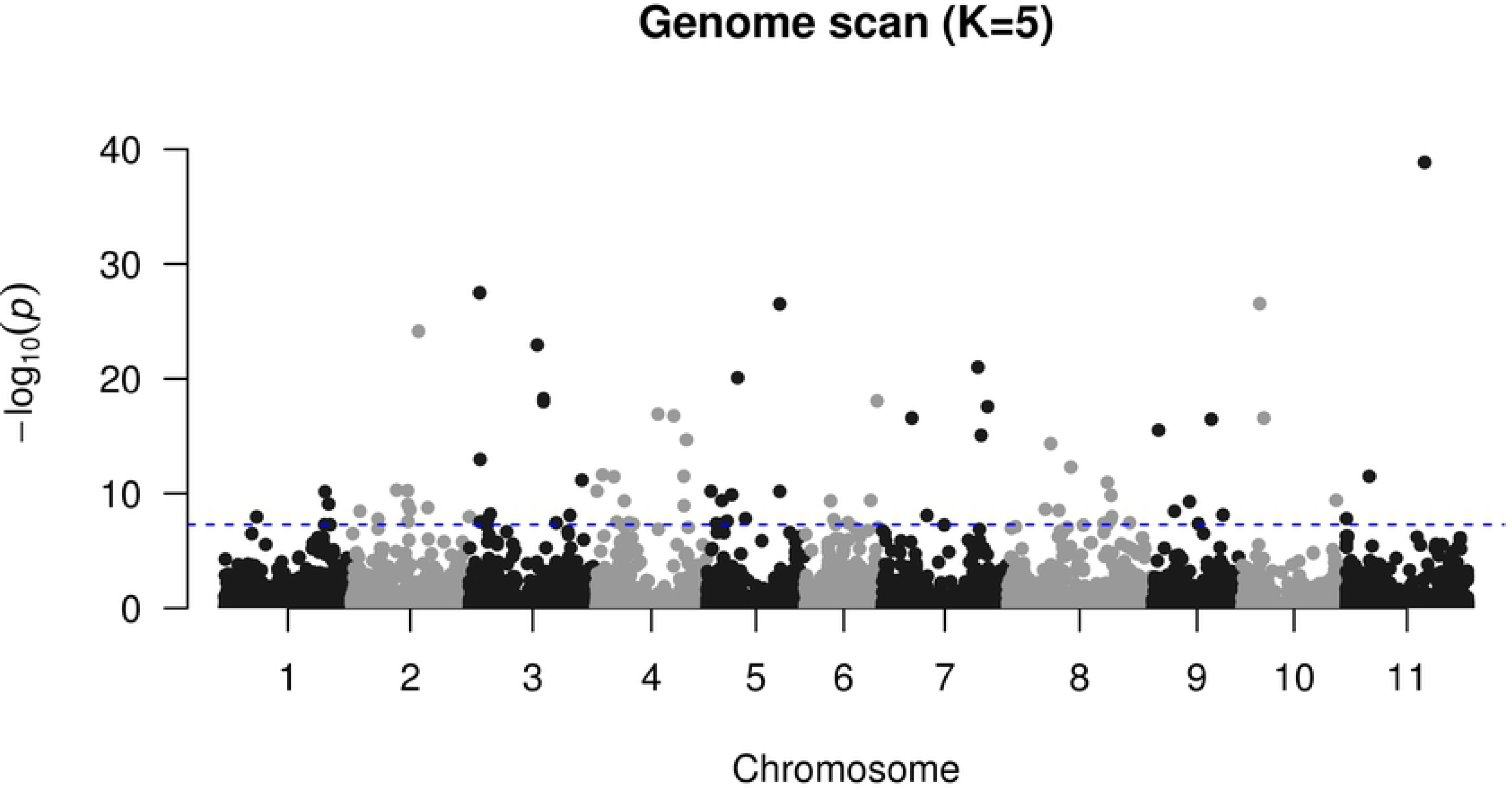
Manhattan plot of the genome scan data with 5 sub-populations (K). The blue dashed line represents the significance threshold (Bonferroni p-value ≤ 0.001).

Of the 70 genes identified by the analysis, 20 of them (28%) were located within a haplotype block (**S1 Table**). When mapping the significant SNPs identified by genome scan analysis to the latest reference genome (v2.1), 62 genes (88%) were confirmed as putatively under selection also in this assembly (**S1 Table**). Interestingly, among the genes not identified in v2.1, three were not present in the annotation file, while one gene (Phvul.001G080400) was renamed Phvul.L006501 and was located to a scaffold instead of chromosome 1.

Among the genes identified, we found several related to drought and/or abscisic acid (ABA) response. Phvul.002G331700, a homolog of the *Arabidopsis* KUP6, is involved in potassium uptake transporter and stomata movement and Phvul.002G143100 is a glycine-rich domain protein (GRP) involved in auxin signaling and stress response. Phvul.004G102800 is a homolog to *Arabidopsis* SLAH3 involved in ABA response; Phvul.008G161000 is a homolog of *Arabidopsis* CAO, a gene related to chlorophyll biosynthesis and ABA signaling; and Phvul.009G050600 is a gene annotated as an importin β protein involved in ABA and drought response in *Arabidopsis*.

### Genome-wide association analysis

A genome-wide association analysis identified 49 genes associated with the bio-climatic variables selected for this analysis. Except for the bio_18 variable (Precipitation of Warmest Quarter), for which no associations were detected, the other variables were associated with at least one gene. The bio-climatic variables with the highest number of associated genes were bio_3 (Isothermality) with 29 genes, and bio_12 (Annual precipitation) with 11 genes (**S2 Table**).

The associated genes were located in all 11 common-bean reference genome chromosomes, except for chromosome Pv06 where there were no significantly associated SNPs. Some of the genes were associated with more than one bio-climatic variables (Fig 5), suggesting the possibility that they could be related to multiple environmental stimuli.

**Fig 5.**
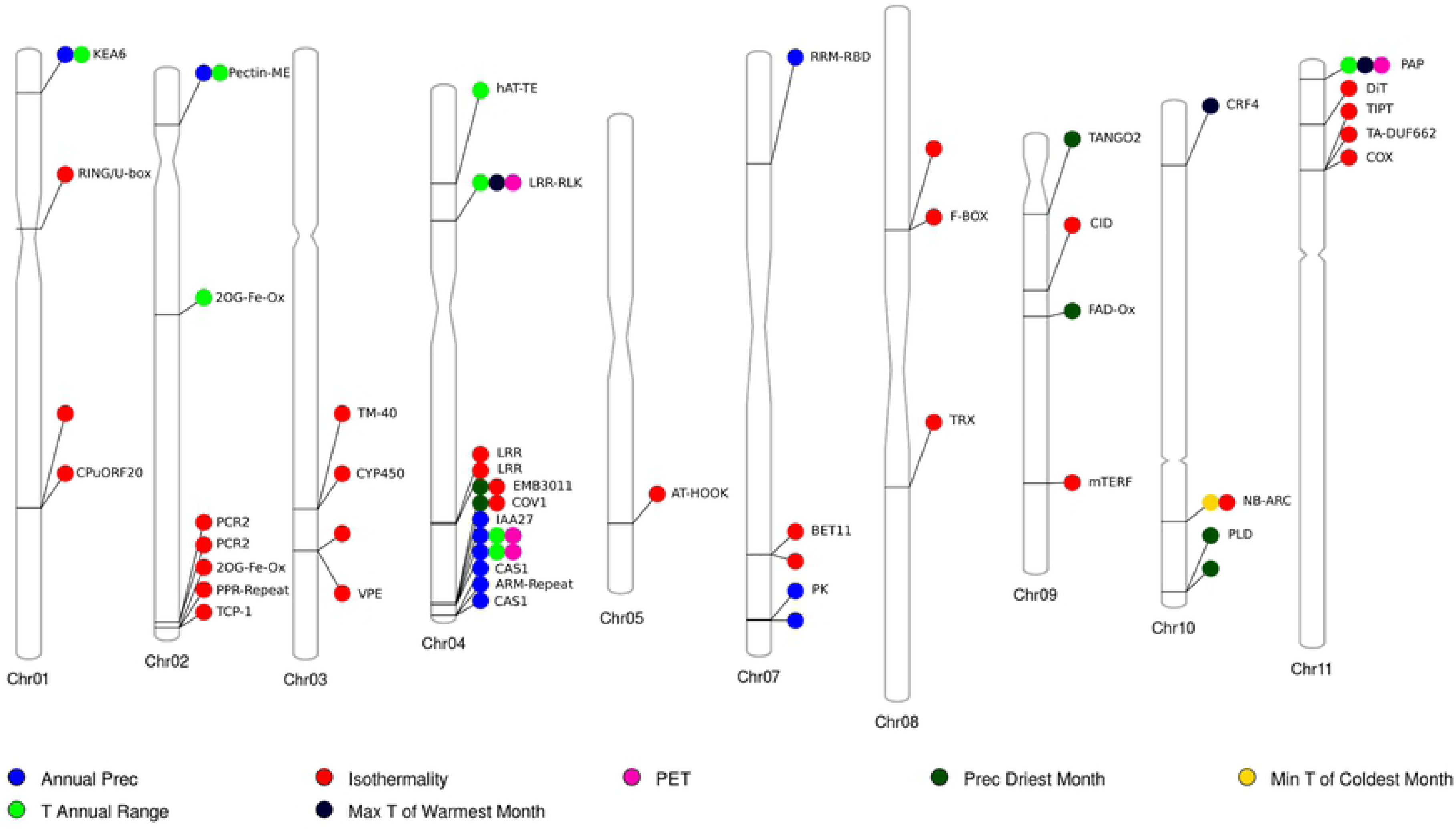
Chromosome ideogram of the genes identified as associated with the bio-climatic variables analyzed. Only chromosomes with significantly associated variants are shown. Each circle represents a different bio-climatic variable. When available, gene annotations are shown. The centromeric regions shown are based on the results from Sevilla et al. (2015).

Of these 49 genes identified by genome-wide association analysis, only 10 (20%) were located within a haplotype block (**S2 Table**). In addition, when mapping significant SNPs identified by association analysis to the latest reference genome (v2.1), 44 genes (88%) were confirmed as putatively associated with environmental variables also in this assembly (**S2 Table**). Four out of five of the missing genes were not present in the v2.1 annotation file.

Among the genes significantly associated with one or more bio-climatic variables, we found several of them related to hormone response, ion homeostasis, plant development, metabolism, and response to stress, in particular drought (**S2 Table**). Among the genes identified, we found some interesting candidates probably involved in stress resistance, like Phvul.001G034400, a homolog of *Arabidopsis* KEA6 involved in potassium homeostasis; Phvul.010G155000, homologous to an *Arabidopsis* phospholipase D α 1 (PLDα1) involved in ABA signaling; Phvul.010G035200 homolog of a cytokinin responsive factor homologous of *Arabidopsis*; and Phvul.008G161700, homologous to an *Arabidopsis* thioredoxin involved in ROS signaling. Interestingly, there was no overlap between the genes identified by genome scan and association analysis.

### Candidate gene allele distributions

To evaluate the geographic distribution of alleles in candidate genes identified by genome scan and association analysis, we clustered the genotypes into groups with a K-means clustering approach on the molecular PCs calculated with pcadapt. The advantage of a K-means clustering approach, over a standard population structure analysis, is that it clearly assigns individuals to specific clusters. The K-means clustering approach identified three clusters for the MW group, with two clusters (MW1 and MW2) located in Mexico and another (MW3) in Central America and Colombia, plus one cluster each for the intermediate (PhI) and the Andean (AW) group (**S4 Fig**). Interestingly, the clustering results closely resembled those obtained in a previous study with more advanced population structure approaches (**S3 Table**) (Ariani et al., 2018).

The allele frequency distribution of the candidate genes identified by genome scan showed drastic differentiation between the genetic groups identified (Fig 6), as expected from the assumptions of the genome scan approach, with some alleles being private for just one of the genetic group (like the alternative alleles for GRP and CAO that were observed only in the AW group). On the other hand, the genes identified by association analysis showed a wide variety of allele frequencies distribution across the different genetic groups (Fig 7), even though some genes had only a single allele in some of the populations (like the reference allele for PLD and TRX in the PhI and AW group). In general, the genes identified by association analysis showed a higher variation of allele frequencies among the different MW groups.

**Fig 6.**
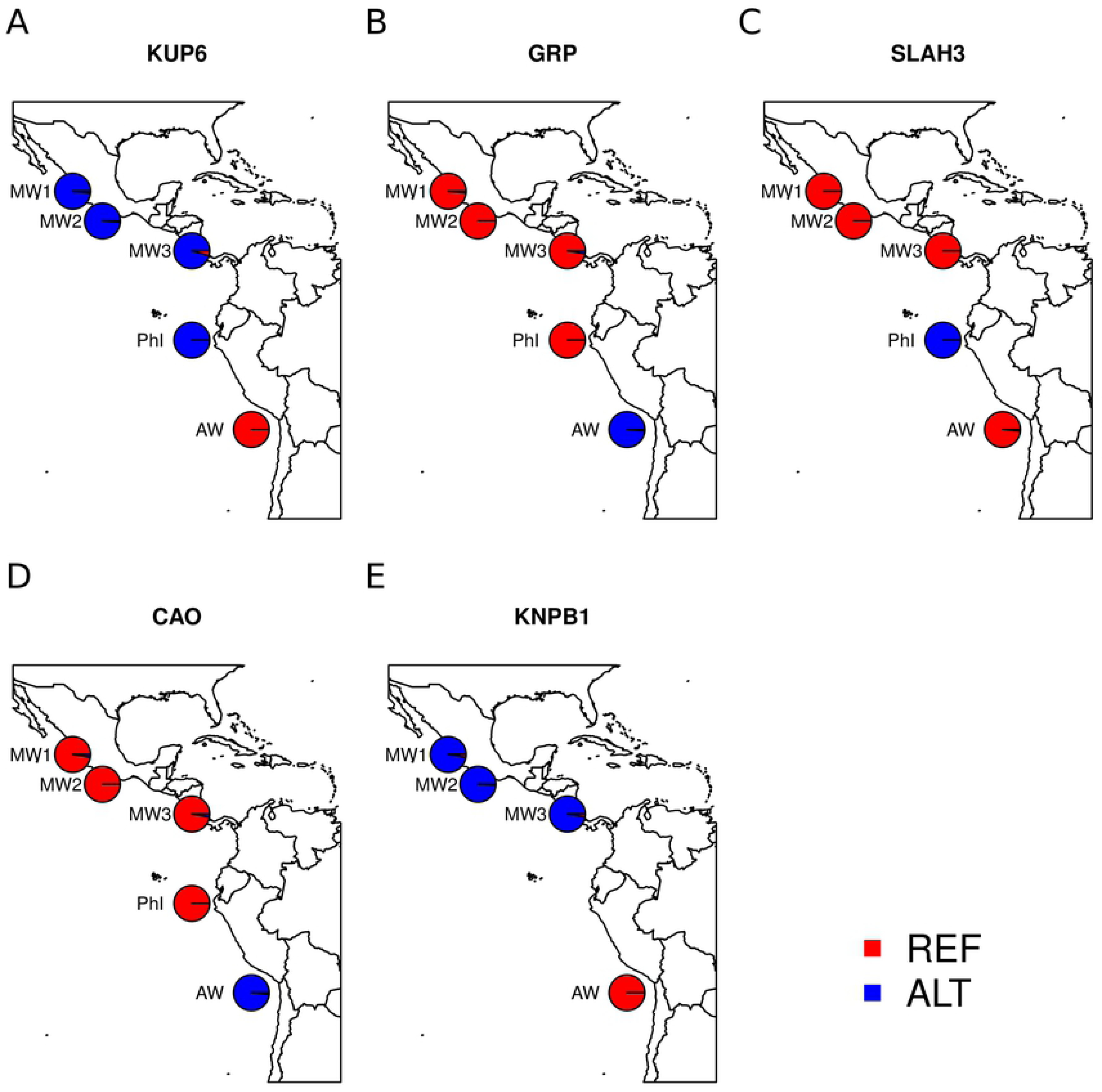
Allele frequency distribution across different genetic groups for candidate genes identified by genome scan analysis. *P. vulgaris v1.0* genes annotation and ID: **(A)** Potassium uptake transporter (Phvul.002G331700); **(B)** Glycine-rich domain protein (Phvul.002G143100); **(C)** ABA response (Phvul.004G102800); **(D)** Chlorophyll biosynthesis and ABA signaling (Phvul.008G161000); **(E)** ABA and drought response (Phvul.009G050600). For panel **(E)** the PhI group was removed because SNP data were completely missing. REF: Reference allele, in red, ALT: Alternative allele, according to the *P. vulgaris* v1.0 gene version, in blue (Sevilla et al., 2015).

**Fig 7.**
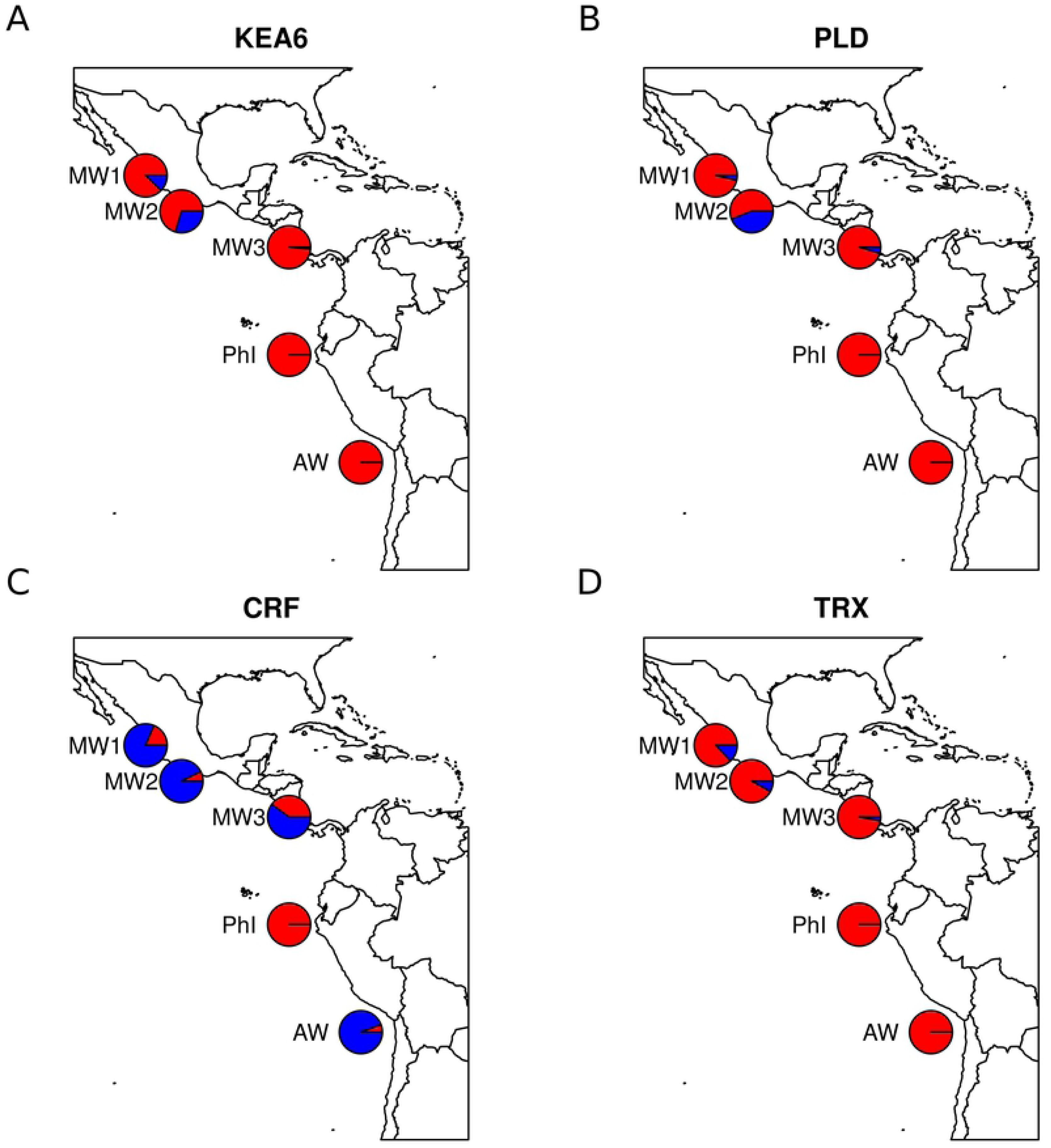
Allele frequency distribution across different genetic groups for candidate genes identified by association analysis. *P. vulgaris v1.0* genes annotation and ID: **(A)** Potassium efflux antiporter (Phvul.001G034400); **(B)** Phospholipase D α 1 (Phvul.010G155000); **(C)** Cytokinin responsive factor (Phvul.010G035200); **(D)** Thioredoxin (Phvul.008G161700). Reference and Alternative alleles are colored as in Fig 6.

## Discussion

Wild common bean (*P. vulgaris*) grows in several areas of Mexico and Central and South America, from northern Mexico to northwestern Argentina across ∼70 latitudinal degrees, in different environments with a wide range of altitudes, average temperatures, and rainfall regimes (Cortés et al., 2013, Gepts 1998, Porch et al., 2013). Thanks to this exceptional geographic distribution, its complex evolutionary history, and high levels of genetic diversity, this species represents an extraordinary resource for evolutionary studies (Chacón et al., 2007, Koenig and Gepts, 1989, Mamidi et al., 2013, Rendón-Anaya et al., 2017a,b, Bitocchi et al., 2012, Kwak and Gepts, 2009), but can be also a conceptual framework for testing and validating landscape genomics approaches in wild plant populations and its feasibility for breeding improvement of domesticated crops (Anderson et al., 2016). In the current study, we identified several genes that could be involved in environmental adaptation in wild common bean by combining genome scan and association analysis. If validated, the genes identified could be useful candidates for improving stress resistance in domesticated common bean. The concordance between the genes tagged by significant SNPs identified between the two different reference genomes assembly suggests also high concordance between the two versions, with possible minimal differences due to the different sequencing data used for the assembly. In addition, the integration of haploblocks information with the genes identified by our analysis showed that most of these genes (70-80%) are located in regions with low LD in wild common bean. This result suggests that those same genes could be located in regions with relatively high recombination frequencies, thus facilitating possible introgression into the domesticated gene pool.

### Genome scan of selection

Molecular PCA analysis clearly separated the three major groups of this species (MW1-MW3, PhI, and AW), as observed in previous research. In particular, the Intermediate gene pool (PhI) was shown again to most diverged group from the Mesoamerican and Andean gene pools, especially along the molPC2 and molPC3 axes (Fig 2), further supporting the hypothesis that this gene pool is actually a distinct species of *Phaseolus* (Rendón-Anaya et al., 2017a,b). A genome scan based on molecular PCA analysis identified several genes with a strong signature of selection (hard-selection sweep) that could be involved in environmental adaptation across the geographical range of this species. The identification of several genes involved in plant development and hormone and stress response, suggests that the different populations of this species adapted to their environment by integrating and adjusting to developmental, hormonal and environmental cues. Several genes among those identified could reflect adaptation to abiotic stress. These genes are also of interest for improving stress resilience in common bean, like the KUP6 potassium (K^+^) transporter located on chromosome Pv02 (Phvul.002G331700). This gene has been directly linked to drought stress by regulating ABA response and stomata movements in *Arabidopsis* (Osakabe et al., 2013). A homolog of this gene located on chromosome Pv03 (Phvul.003G052900) showed a higher genetic and transcriptional diversity in Mesoamerican domesticated beans than in wild ones (Bellucci et al., 2014; Bitocchi et al., 2017). Due to the possible role of KUP-like genes in response to drought stress and their identification as selected genes in both wild and domesticated populations of common bean, further studies should focus on the evolution and diversity of this gene family in this species.

Another gene identified in the current study and possibly involved in adaptation to drought response in wild common bean is Phvul.004G102800, homolog of SLAH3 of *Arabidopsis*, which was annotated as an S-type anion channel. This type of channels is rapidly regulated by ABA and stimulates stomata closure by inhibiting inward K^+^ channels, thus reducing K^+^ influx into guard cells (Geiger et al., 2011; Zhang et al., 2016). In addition to being involved in drought stress response, this same gene has been recently identified also as related to salinity stress response in *Arabidopsis* by regulating ion homeostasis between root and shoots (Cubero-Font et al., 2016).

The chlorophyll alpha oxygenase (Phvul.008G161000), identified as a gene under selection (and a homolog of *Arabidopsis* CAO), has a primary role in the biosynthesis of chlorophyll b (Espineda et al., 1999). However, *Arabidopsis* mutants for this gene showed a reduction of antioxidant compounds (specifically glutathione) in guard cells and an increased ABA sensitivity in comparison to wild type plants (Jahan et al., 2016), suggesting a possible involvement of this gene in adaptive response to stressful environments.

Phvul.002G143100, identified as selected within the different sub-populations of *P. vulgaris*, is annotated as a glycine-rich domain protein, homologous of *Arabidopsis* GRDP2 gene. GRP are a multi-gene superfamily present in several organisms, including plants (Sachetoo-Martins et al., 2000). This gene family has been associated in plants with several developmental processes and in responses to both biotic and abiotic stresses (Mangeon et al., 2010). A recent study focusing on the characterization of the direct *Arabidopsis* homologs of this gene (ATGRDP2) demonstrated that this gene regulates plant growth and flowering by accumulating higher level of indole-3-acetic acid and improves abiotic stress response (Ortega-Amaro et al., 2014). In particular, the over-expression of this gene in transgenic plants increased growth rate and reduced days to flowering. It also increased salt tolerance in comparison to wild-type plants.

In addition to the previous genes identified as selected by genome scan analysis and putatively involved in environmental response in plants, Phvul.009G050600 was identified and annotated as an importin β-protein homologous to *Arabidopsis* KPNB1. This gene mediates the import of proteins and protein complexes between the cytoplasm and the nucleus and is essential in regulating signal transduction pathways in response to environmental and developmental stimuli (Merkle, 2003). In particular, the *Arabidopsis* homolog of Phvul.009G050600 (AtKPNB1) has been directly related to ABA and drought response previously (Luo et al., 2013).

### Genome-wide association analysis

Association analysis between genotypic data and bio-climatic variables identified several genes significantly associated with one or more bio-climatic variables, putatively involved in plant development, ion homeostasis, and stress response. Among these genes, several could be useful as potential molecular markers for improving abiotic stress in domesticated common bean. As examples, we identified a gene related to potassium homeostasis and annotated as a K^+^ efflux antiporter (KEA) gene associated with bio_12 (Annual Precipitation) and bio_7 (Temperature Annual Range). Potassium is an essential macronutrient involved in several physiological and developmental processes in all living organism, and in plants this cation is also essential in maintaining plant osmotic potential, cytosolic pH, and stomata movement (Shabala, 2003, Sharma et al., 2013). In addition, variation in K^+^ homeostasis is one of the first responses to several abiotic and biotic stresses in plants, allowing the plants to rapidly respond to stressful conditions (Shabala and Pottosin, 2014) and making the KEA gene identified in the current study an interesting candidate gene for further analysis.

Another gene, significantly associated with bio_14 (Precipitation of Driest Month) is Phvul.010G155000, which is annotated as a phospholipase (PLDα1). This gene is involved in the biosynthesis of phosphatidic acid (PA), which is an important signaling molecule in response to several stresses in plants (Saucedo-García et al., 2015). In particular, PA is involved in the ABA signaling cascade and regulates stomata closure in plants by directly interacting and blocking ABI1, an inhibitor of ABA response in plants (Zhang et al., 2004). This gene regulates stomatal closure and ABA-dependent hydrogen peroxide (H_2_O_2_) production in *Vicia faba* as well (Qu et al., 2014), making this gene an interesting candidate for improving drought response in common bean.

An additional gene, significantly associated with bio_5 (Max Temperature of Warmest Month), is Phvul.010G035200, annotated as a cytokinin response factor homologous of *Arabidopsis* CRF4. Cytokinin is an essential plant hormone involved in growth and developmental processes (Durán-Medina et al., 2017, Kieber and Schaller, 2014), but in recent years it has also been implicated in the response and adaptation to different environmental stresses (Novakova et al., 2007, O'Brien and Benková, 2013). CRF genes are a class of plant transcription factors responsive to cytokinin that integrate hormonal and environmental signals for adapting plant growth and development in response to the environment (Kim, 2016, Rashotte and Goertzen, 2010). The *Arabidopsis* homolog of this gene has been previous related to acclimation to cold temperatures (Zwack et al., 2016). Since this gene has been associated with temperature variables in wild common bean, it could also be involved in adaptation to temperature variation in this species.

Another gene of interest, Phvul.008G161700, is significantly associated with bio_3 (Isothermality) and is annotated as a thioredoxin protein. These proteins are involved in the regulation of oxidative stress response and in scavenging reactive oxygen species (ROS) in plants (Gelhaye et al., 2005). Other than being simple byproducts of cellular metabolism, ROS molecules has been recognized as important signaling molecules that regulate the response to several environmental stresses in plants (D’Autréaux and Toledano, 2007, Sewelam et al., 2016). Due to their ability to control the redox state of the cell, thioredoxin represents a key component of the ROS signal transduction pathways in plants and in the response to environmental stress (Sevilla et al., 2015). Thus, this gene could constitute another interesting candidate gene for improving stress resistance in domesticated common bean.

### Comparison of genes identified by genome scan and GWAS

Even though the genes identified by outlier-detection methods (hard-selection sweeps) and association methods (soft-selection sweeps) are involved in similar processes, there was no overlap between the candidate genes identified by the two approaches in this study. This could be the direct result of the different assumptions underlying these methods. Indeed, genome scan analysis identify genes that shows drastic variations of allele frequencies between natural subpopulations (Schoville et al., 2012, Wagner and Fortin, 2013). This approach is independent from bio-climatic variables, thus the SNPs identified as under selection by this analysis could be the results of selective mechanisms not considered by association analysis, like soil composition, pathogen pressure and/or competition with other plants. On the other hand, association analyses identify SNPs showing slight variations in allele frequencies across environmental gradients that can increase environmental adaptation in natural populations (Schoville et al., 2012, Wagner and Fortin, 2013). This selection process usually acts on natural standing variations and favor the presence of multiple alleles and haplotypes, instead of allele fixation within populations (Hermisson and Pennings, 2005).

### Epilogue

In conclusion, landscape genomic analysis of wild common bean genotypes allowed us to identify several genes showing a signature of presumed selection in this species. It is likely that two methods – genome scan and GWAS - are indeed complementary for understanding local adaptation in wild plant populations, as observed previously in other species (Dell'Acqua et al., 2014, Pyhäjärvi et al., 2013) and are a feasible approach for the preliminary identification of novel candidate genes for adaptation to climatic differences along the exceptionally broad habitat of wild common bean. Further corroboration of the actual role of the candidate genes in adaptation will come from introgression of these genes from wild to domesticated beans and a concurrent phenotypic analysis showing improved performance under stress conditions.

Our long-term objective is to identify both populations (Ariani et al., 2018) and genes (this study) that have been putatively under selection by the abiotic stresses of temperature, rainfall, and the related variable, potential evapotranspiration. Identification of these potential sources of genetic tolerance are only a first step towards the development of more stress-resilient beans. The next step is to corroborate the effectiveness of these wild populations and these candidate genes as a source of stress-tolerance through indirect selection for these genes in selected populations resulting from the cross between candidate populations and domesticated testers (Acosta-Gallegos et al., 2007). In a recent paper, Cortés and Blair (2018) identified 115 SNPs tagging 77 annotated genes potentially selected by drought tolerance among a set of wild bean populations, using correlations between SNPs and an average yearly drought index. They argued that drought tolerance and performance under well-watered conditions were mutually incompatible. The testcrosses just mentioned between domesticated testers and wild populations that have been subjected to different drought stress conditions, as identified in this study and Ariani et al., (2018), will allow us to examine this hypothesis, which has considerable implications for breeding for stress tolerance.

## MATERIALS AND METHODS

### Plant material and genotypic data

A panel of 246 wild *P. vulgaris* accessions, previously genotyped with a Genotyping-By-Sequencing (GBS) protocol using the *Cvi*AII restriction enzyme (Ariani et al., 2018), was analyzed. The panel was representative of the ecological and geographic distribution of this species and included 157 genotypes of the Mesoamerican (MW), 77 of the Southern Andes (AW), and 12 of the Central Andes (Northern Peru-Ecuador; PhI) gene pools. The SNPs considered in this study were those with a Minor Allele Frequency (MAF) ≥ 0.05 and less than 20% missing data. The list of the accessions sequenced, with gene pool information and geographic coordinates, is available in **S4 Table**, while genotyping data in VCF format are available as a Dash dataset (https://doi.org/10.25338/B8DW39). The seeds were provided by the Genetic Resources Unite at the International Center of Tropical Agriculture (CIAT, Cali, Colombia) and the United States Department of Agriculture Western Regional Plant Introduction Station (Pullman, WA).

### Spatial Analysis

Spatial analyses were conducted within the R statistical environment (www.r-project.org) using the dismo package and its dependencies (raster and sp). The geographic coordinates of the individuals analyzed in this study were used for retrieving the 19 bio-climatic summary variables from the WorldClim database (http://www.worldclim.org/). The data were downloaded at a 30-second resolution (approximately 0.86 km^2^ at the equator). In order to identify a subset of bio-climatic variables that best summarizes our dataset, we performed a Principal Component Analysis (PCA) on the scaled and centered variables using the ChemometricsWithR package (Wehrens 2011). We then selected the first two variables with the highest positive and negative loading in the first four principal components (PC1 to PC4) (**S3 Table**). Since some of the selected bio-climatic variables showed a high correlation (**S1 Table**), we decided to pick only one of the correlated variables for further analysis. The final bio-climatic variables analyzed in this study were: bio_3 (Isothermality), bio_5 (Max Temperature of Warmest Month), bio_6 (Minimum Temperature of Coldest Month), bio_7 (Temperature Annual Range), bio_12 (Annual precipitation), bio_14 (Precipitation of Driest Month), and bio_18 (Precipitation of Warmest Quarter). In addition to the above-mentioned bio-climatic variables, we included also annual Potential EvapoTranspiration (PET) downloaded from the Global Aridity and PET Database (http://www.cgiar-csi.org/data/global-aridity-and-pet-database).

### Genome Scans for Selection and Association Analysis

Genome scans for selection (i.e., hard selective sweeps) were performed on the final set of SNPs using the pcadapt R package (Luu et al., 2017), an algorithm able to detect population structure and outlier loci by performing a PCA analysis on SNP genotypic data. The best number of sub-populations was inferred by visually evaluating the scree plot of eigenvalues for the different principal components (K); the genomic scans for selection were performed for K in the range 2-5. The p-values obtained by this analysis were corrected using the Bonferroni method and only SNPs with a corrected p-value ≤ 0.001 were considered as significant.

Association analysis (i.e., soft selective sweeps) was performed separately for each of the seven selected bio-climatic variables and annual PET. For this analysis, we used the LFMM algorithm (Frichot et al., 2013) implemented in the LEA R package (Frichot and François, 2015). This method was developed specifically for identifying signature of environmental selection in genomic data and can efficiently correct for population history and isolation-by-distance (IBD). To correct for spurious association determined by population structure or IBD, the number of latent factors (i.e., populations) needs to be decided *a priori* and subsequently evaluated using the genomic inflation factor parameter. Since LFMM is based on Monte Carlo Markov Chain (MCMC) sampling, we ran it multiple times for each association analysis and then averaged the p-values (as suggested in the software documentation). To identify the best number of populations (K) for association with each bio-climatic variable, we performed three runs of the program with K in the range 4-10 and estimated the inflation factor from these runs (Devlin and Roeder, 1999). Plots of the inflation factor for different values of K (**S5 Fig**) showed that the best inflation factor for reducing False Discovery Rate (FDR) (i.e., closest to 1) was six for Bio12, Bio14, and Bio5, and 7 for Bio6, Bio18, Bio7, Bio3, and PET. Based on this preliminary screening, we re-ran the program with the best number of K for 10 times with 10,000 MCMC iterations and a burn-in period of 1,000. The p-values where then averaged across the different runs and corrected using the Bonferroni method. SNPs with a corrected p-value ≤ 0.05 were considered as significant.

### Identification of putatively selected genes

The distance between significant SNPs, identified by genome scans or association analysis based on the *P. vulgaris* v1.0 genome annotation (https://phytozome.jgi.doe.gov/pz/portal.html) (Schmutz et al., 2014), was evaluated using the GenomicRanges/rtracklayer packages or R (Lawrence et al., 2009, 2013). Only genes within 5 Kb of a significant SNPs were chosen as putatively selected genes. This 5 Kb upper limit was selected based on the genotyping approach used in this study (that did not allow a full coverage of the genome), but also considered the presence of possible regulatory regions immediately adjacent to gene sequences (Li et al., 2012). To understanding if the genes identified by significant SNPs were in regions with high linkage disequilibrium (LD), we identified haploblocks from the complete set of SNPs data using the PLINK program (Purcell et al., 2007) with default parameters. For downstream analysis, we considered only blocks longer than 100 bp. We then integrated this information with the genes identified as putatively selected by genome scan or association analysis, to determine if these candidate genes were located in haploblock regions. This analysis identified 1338 haplotype blocks evenly distributed across the 11 chromosomes (Dash dataset: https://doi.org/10.25338/B8DW39).

### Comparison with latest genome reference

A new genome reference for *P. vulgaris* (v2.1) has been released on Phytozome although it has yet to be peer-reviewed. We compared the genes and the SNPs identified by our analysis between the old (v1.0) and the newest (v2.1) genome version. To compare the results between the two genomes, we mapped the significant SNPs, identified by genome scan and association analysis in the v1.0 genome reference, onto the v2.1 version. For this analysis, we extracted the 100 bp upstream and downstream of a significant SNPs (200 bp window) in the v1.0 version and mapped them to the v2.1 reference genome using nucleotide BLAST (Camacho et al., 2009). For each SNP and relative flanking region, we then selected the best hit in the v2.1 genome and identified the genes annotated in the new reference located within 5 Kb of the hit (as described in the ‘Identification of putatively selected genes’ section).

### Candidate genes evaluation across genetic groups

For clustering individuals based on genetic groups and visualizing allele frequency variations across clusters, we applied a K-means clustering approach using the first 5 PCs obtained from pcadapt analysis. We selected K=5 as the best number of clusters, based on the scree plot of the eigenvalues obtained with pcadapt. The clustering analysis was performed using the python scikit-learn library (Pedregosa et al., 2011). For each genetic cluster, we calculated allele frequencies for SNPs tagging candidate genes using VCFtools (Danecek et al., 2011) and plotted them on genetic maps using R.

## Acknowledgements

This work used the Vincent J. Coates Genomics Sequencing Laboratory at UC Berkeley, supported by NIH S10 Instrumentation Grants S10RR029668 and S10RR027303. We thank the Genetic Resources Unit of the Centro Internacional de Agricultura Tropical (CIAT, Cali, Colombia) and the Western Regional Plant Introduction Station of the USDA (Pullman, WA) for providing samples of wild *P. vulgaris* used in this study.

## Availability of Data and Materials

Raw sequencing data are available at the NCBI Sequence Read Archive (http://www.ncbi.nlm.nih.gov/sra) under the accession numbers SRX2771627 and SRX2771628. The variants file and the relative geographical coordinates used for performing the analysis are available as additional files of the current manuscript.

## Authors’ contributions

AA performed the experiment, analyzed the data and wrote the manuscript. PG designed the experiment, supervised the work and wrote the manuscript. All authors approved the final version of the manuscript.

## Ethics approval and consent to participate

Not applicable

## Consent for publication

Not applicable

## Supporting information

### Tables

**S1 Table.** Candidate genes identified by genome scans.

**S2 Table.** Candidate genes identified by genome-wide association analysis.

**S3 Table.** Eigenvalues of the different bioclimatic variables along the first four principal components.

**S4 Table.** List of the final wild *Phaseolus vulgaris* analyzed in this study. Accession ID, country of origin, geographical coordinates of collection, and gene pool information are shown (from Ariani et al. 2018).

### Figures

**S1 Fig.** Correlation graphs between bio-climatic variables for the different *P. vulgaris* accessions analyzed. Correlation coefficients are rendered using circles (upper-right part) or by showing directly the value (lower-left part). Color are based on color-bar in the right side of the graph.

**S2 Fig.** Cumulative variance explained by the different PCs when performing a PCA on bio-climatic variables.

**S3 Fig.** P-values distribution for genome scans with 2 (A), 3 (B), 4 (C) or 5 (D) sub-populations.

**S4 Fig.** Plot of geographic distribution of the wild *P. vulgaris* analyzed in the current studies. Genotypes are colored based on the different clusters identified by K-means clustering (Fig 1B, Fig. 2B, **S4 Table**).

**S5 Fig.** Plots of the inflation factor for different values of K across the climatic variables selected for association study.

